# Uncovering the microbiome of invasive sympatric European brown hares and European rabbits in Australia

**DOI:** 10.1101/832477

**Authors:** Somasundhari Shanmuganandam, Yiheng Hu, Tanja Strive, Benjamin Schwessinger, Robyn N. Hall

## Abstract

European brown hares (*Lepus europaeus*) and European rabbits (*Oryctolagus cuniculus*) are invasive pest species in Australia, with rabbits having a substantially larger environmental impact than hares. As their spatial distribution in Australia partially overlaps, we conducted a comparative microbiome study to determine how the composition of the gastrointestinal microbiota varies between these species, since this may indicate species differences in diet, physiology, and other internal and external factors. We analysed the faecal microbiome of wild hares and rabbits from a sympatric environment, additionally comparing Illumina and Nanopore sequencing platforms. The faecal microbiomes varied significantly between hares and rabbits, despite both species occupying a similar habitat. Moreover, we identified significantly more variation in faecal microbiome composition between individual rabbits compared to hares. The faecal microbiome in both species was dominated by the phyla *Firmicutes* and *Bacteroidetes*, typical of many vertebrates. Many phyla, including *Actinobacteria*, *Proteobacteria*, and *Patescibacteria*, were shared between rabbits and hares. In contrast, bacteria from phylum *Verrucomicrobia* were present only in rabbits, while phyla *Lentisphaerae* and *Synergistetes* were represented only in hares. We did not identify phylum *Spirochetes* in Australian hares; this phylum was previously shown to be present at high relative abundance in European hare faecal samples. These differences in the faecal microbiota between hares and rabbits in Australia may be associated with differences in diet, and potentially behaviour, of the host species in their non-native range, which may influence the environmental impacts that these species have in Australia.

## Introduction

In a new environment, non-native species must face several barriers to first invade and then become established. They must quickly adapt to new environmental conditions while also competing with native species for food, shelter, and water (1, 2). European rabbits (*Oryctolagus cuniculus*) and European brown hares (*Lepus europaeus*) are lagomorphs from the family *Leporidae* and are both non-native species in Australia, originally introduced from Europe in 1859 (3). European rabbits rapidly colonised the entire continent and are considered to be one of Australia’s worst invasive vertebrate pests due to land degradation and competition with native animals and livestock (4–6). Despite foraging over wider areas, the impacts of European brown hares are less severe, although they are still considered to be a pest species (6, 7). The specific factors underlying differences in colonising potential and impacts of each species are likely multifactorial, including differences in host physiology, reproductive strategies, behaviour, diet, and interaction with commensal and pathogenic microbes.

Recent advancements in sequencing technologies have highlighted the importance of interactions between the gut microbiome, host behaviour and physiology, and dietary preferences (8–13). Although the gastrointestinal microbiome of both rabbits and hares have previously been investigated, most studies to date were conducted on domestic rabbit populations in either Europe or China (14–24). Evaluating the microbial diversity in the gastrointestinal tract of wild hares and rabbits in Australia may provide additional insights as to why European rabbits were such successful colonisers in this environment compared to hares (25). Furthermore, identifying differences between the gastrointestinal microbiota of lagomorphs in their native range compared to their introduced range may lead to novel insights into how to more sustainably manage introduced populations (25).

The microbiome can be investigated through a range of both traditional culture-dependent and less biased culture-independent techniques, including high throughput sequencing (26–31). Among various sequencing strategies, taxonomic profiling based on the 16S ribosomal RNA (rRNA) gene is widely used for estimating the bacterial and archaeal diversity in tissue or environmental samples due to the availability of well-curated databases such as SILVA (32). Broadly, high-throughput sequencing can be classified as either short-read sequencing, using the Illumina platform, or long-read sequencing, using platforms such as PacBio, IonTorrent, or Oxford Nanopore (33, 34). Although all these platforms have previously been used to conduct 16S rRNA microbial profiling studies, direct comparisons of these methods are limited (35).

To investigate the differences between the gastrointestinal microbiome of Australian wild rabbits and hares we conducted 16S rRNA sequencing on faecal samples collected from sympatric populations of these species using both Illumina and Oxford Nanopore sequencing platforms. The gastrointestinal microbial diversity of Australian lagomorphs was then compared to that observed in European populations to explore differences between populations in their native and invasive ranges.

## Materials and Methods

### Sample collection

Faecal samples were collected at necropsy from nine wild European brown hares (*Lepus europaeus*) and twelve wild European rabbits (*Oryctolagus cuniculus*) of both sexes (Table S1). The hares and rabbits used in this study were mostly adults or young adults, they were outwardly healthy, and were shot as part of routine vertebrate pest control operations in Mulligans Flat, ACT (-35.164, 149.165), between January and September 2016. Mulligans Flat Nature Reserve comprises 781 hectares of box-gum grassy woodland with a sparse to moderately dense ground cover of grasses, herbs, and shrubs. The reserve is fenced, preventing immigration and emigration of most animals. Rabbits and hares were shot from a vehicle using a 0.22-caliber rifle targeting the head or chest. Faecal pellets from the descending colon were collected during post-mortem and stored at -20 °C. All sampling was conducted according to the Australian Code for the Care and Use of Animals for Scientific Purposes as approved by CSIRO Wildlife and Large Animal Ethics Committee (approvals #12-15 and #16-02).

### Genomic DNA extraction

Genomic DNA was extracted from 50 mg of faecal pellets using the DNeasy Blood & Tissue kit as per manufacturer’s instructions and described in detail at https://www.protocols.io/view/genomic-dna-extraction-from-animal-faecal-tissues-6bbhain (Qiagen, Chadstone Centre, Victoria). Hare and rabbit samples were processed separately, and reagent-only controls (ROC) were included with each set of extractions. All samples were quantified and evaluated for integrity using the Qubit dsDNA Broad Range assay kit and the Qubit Fluorometer v2.0 (Thermo Fisher Scientific, Waltham, MA) and Nanodrop ND-1000 (Thermo Fisher Scientific). Genomic DNA samples were stored at –20 °C.

### Illumina library preparation, sequencing, and bioinformatics analysis

We amplified the V3-V4 region of the 16S rRNA gene (∼ 460 bp) from genomic DNA and ROC according to the Illumina protocol for 16S Metagenomic Sequencing Library Preparation using a dual-indexing strategy, with modifications (36). Briefly, initial PCR reactions (25 μl) were performed for each sample (including ROC) using 2x Platinum SuperFi PCR Master Mix (Thermo Fisher Scientific), 0.5 μM of forward (5’-**TCGTCGGCAGCGTCAGATGTGTATAAGAGACAG**CCTACGGGNGGCWGCAG-3’) and reverse primer (5’-**GTCTCGTGGGCTCGGAGATGTGTATAAGAGACAG**GACTACHVGGGTATCTAATCC -3’) (overhang adapter sequence highlighted in bold) and 4.6 ng of genomic DNA as described at https://www.protocols.io/view/library-preparation-protocol-to-sequence-v3-v4-reg-6i7hchn. A ‘no template control’ (NTC) was included during reaction setup. Cycling conditions were: 98 °C for 30 sec, followed by 25 cycles of 98 °C for 10 sec, 55 °C for 15 sec, and 72 °C for 30 sec, with a final extension of 10 min at 72 °C. PCR products were confirmed by agarose gel electrophoresis before being purified using AMPure XP beads (Beckman Coulter, Indianapolis, IN) at a 1x ratio as described previously (36). Each amplicon was then dual-indexed with unique DNA barcodes using the Nextera XT index kit (N7XX and S5XX, Illumina, San Diego, CA) for PCR-based barcoding (36). For each 50 μl PCR reaction, we used 5 μl of each index (i7 and i5), and 5 μl of the first PCR product. Cycling conditions were as described above but limited to eight cycles. Final libraries (including ROC and NTC) were purified with AMPure XP beads (Beckman Coulter), validated using the Tapestation D1000 high sensitivity assay (Agilent Technologies, Santa Clara, CA) and the Qubit High Sensitivity dsDNA assay kit (Thermo Fisher Scientific), pooled at equimolar concentrations, and sequenced on an Illumina MiSeq using 600-cycle v3 chemistry (300 bp paired-end) at CSIRO, Black Mountain, Canberra.

Illumina fastq reads were analysed in QIIME 2-2019.7 software (37). Raw fastq reads were quality filtered (i.e. filtered, dereplicated, denoised, merged, and assessed for chimaeras) to produce amplicon sequence variants (ASV) using the DADA2 pipeline via QIIME2 (38). The DADA2 generated feature table was filtered to remove ASVs at a frequency less than two, and remaining ASVs were aligned against the SILVA_132 (April, 2018) reference database using QIIME2 feature-classifier BLAST+ (39–41). Fasta sequences from DADA2 were also aligned to the same reference database using BLASTn (2.2.28) through the command line interface (CLI) using an e-value cut-off of 1e-90. We processed and exported BLASTn outputs into QIIME2 to perform taxonomic analysis. All scripts used were deposited at https://github.com/SomaAnand/Hare_rabbit_microbiome. Raw sequence data were deposited in the sequence read archive of NCBI under accession number PRJNA576096.

The microbial diversity and richness between hare and rabbit samples were estimated using alpha diversity (Shannon and observed OTU) and beta diversity (Bray-Curtis dissimilarity metric) metrics after rarefaction at a subsampling depth of 149,523 using the q2-diversity pipeline within QIIME2 (42, 43). This subsampling depth retained all samples except for one ROC (rabbit) and PCR NTC, both of which had low sequence counts.

### Nanopore library preparation, sequencing, and bioinformatics analysis

The entire 16S rRNA gene (∼1500 bp) was targeted for sequencing using the Nanopore MinION sequencing platform. We amplified the 16S rRNA gene from all genomic DNA samples using the universal primers 27F and 1492R [46]. PCR reactions (100 μl) were performed for each sample using 2x Platinum SuperFi PCR master mix (Thermo Fisher Scientific), 0.4 μM each primer, and 11.5 ng genomic DNA as described at https://www.protocols.io/view/library-preparation-protocol-to-sequence-full-leng-6j6hcre. Cycling conditions were: 98 °C for 30 sec, followed by 28 cycles of 98 °C for 10 sec, 55 °C for 15 sec, and 72 °C for 40 sec, with a final extension of 5 min at 72 °C. PCR products were confirmed by agarose gel electrophoresis before being purified with AMPure XP beads (Beckman Coulter), validated using the Tapestation D1000 high sensitivity assay (Agilent Technologies), and quantified using the Qubit High Sensitivity dsDNA assay kit (Thermo Fisher Scientific). For library preparation we used the ligation sequencing kit 1D (SQK-LSK108) in combination with the native barcoding kit 1D (EXP-NBD103) (Oxford Nanopore Technologies, Oxford, UK) as per the manufacturer’s protocol, except 500 ng of input DNA per sample was used for end preparation (44). For the addition of barcodes, 80 ng of end-prepped DNA was used as input, and barcoded samples were pooled in equimolar concentrations to obtain at least 400 ng of pooled DNA. We used a final library amount of 200 ng to obtain maximum pore occupancy on a MinION R.9.4.1 flow cell. Two MinION flowcells were used to run all samples and each flow cell had a mix of hare and rabbit samples to control for potential batch effects.

Nanopore raw reads in fast5 format were demultiplexed using deepbinner (45). Basecalling, adapter trimming, and conversion into fastq format was performed in Guppy 2.3.7 (Oxford Nanopore Technologies). BLASTn was used to locally align 16S sequences with a quality score higher than seven against the SILVA_132 reference database as described above. Top hits were exported in Biological Observation Matrix (BIOM) format and imported into QIIME2 for subsequent analyses. Nanopore data was rarefied at a subsampling depth of 109,435 for diversity analysis. This rarefied dataset was then used to analyse the taxonomic results as per the Illumina workflow described above. Raw sequence data were deposited in the sequence read archive (SRA) of NCBI under accession number PRJNA576096.

### Statistical analysis

We assessed whether the bacterial diversity of hares and rabbits was statistically different by using permutation-based statistical testing (PERMANOVA) via the QIIME diversity beta-group-significance pipeline within QIIME2 for both Illumina and Nanopore datasets (43). Statistical significance in alpha diversity was estimated using the nonparametric Kruskal Wallis test within QIIME2 (46). We also evaluated statistical differences between hare and rabbit faecal samples for each observed bacterial phyla using a combination of multiple tests. We performed analysis of variance (ANOVA) to estimate the variance of both populations and Student’s t-test to identify statistical differences between the means of both groups. Estimated p-values were corrected using the Benjamini-Hochberg method to control the False Discovery Rate for multiple hypothesis testing.

## Results

### Interrogating sympatric hare and rabbit faecal microbiota using 16S rRNA sequencing

We performed Illumina short-read sequencing of the 16S rRNA V3-V4 region and Nanopore long read sequencing of the entire 16S rRNA gene on faecal samples of wild hares and rabbits living sympatrically in a periurban nature reserve in Australia. Short-read sequencing of 24 samples through the Illumina MiSeq pipeline produced an aggregate of 14,559,822 reads greater than Q30 (average 501,908 reads per sample, excluding ROC and NTC). Reads were converted to ASVs, which were assigned using the SILVA_132 reference database via BLAST+ in QIIME2. This produced 6,662 unique features at a frequency greater than two. Long-read sequencing of 21 samples (no ROCs or NTC due to lack of amplification) using the Nanopore MinION platform produced an aggregate of 6,544,770 reads (average of 386,924 reads per sample) above a quality threshold of seven across two MinION runs. After aligning the reads against the SILVA_132 reference database using BLAST, 47,100 unique features with a frequency greater two were obtained.

### Wild rabbits have greater faecal microbial diversity compared to sympatric hares

We conducted alpha and beta diversity analyses to assess the species richness and abundance within and between samples, respectively. Rabbit faecal samples had a significantly higher alpha diversity (p = 0.0001 as measured by Shannon index and Kruskal Wallis test) and beta diversity (p = 0.01 as measured by Bray-Curtis dissimilarity matrix and PERMANOVA) than hare samples. We observed considerable variance in faecal microbial diversity between individual rabbits (Fig. 1). In contrast, the hare faecal samples examined had similar bacterial diversity with relatively low variance between individuals. We observed these effects in both the Illumina and Nanopore 16S sequencing data sets (Fig. 1). We did not observe clear correlation between bacterial diversity and sex, reproductive status, or season of sample collection, although our statistical power was low due to the relatively small sample size.

**Figure 1:**
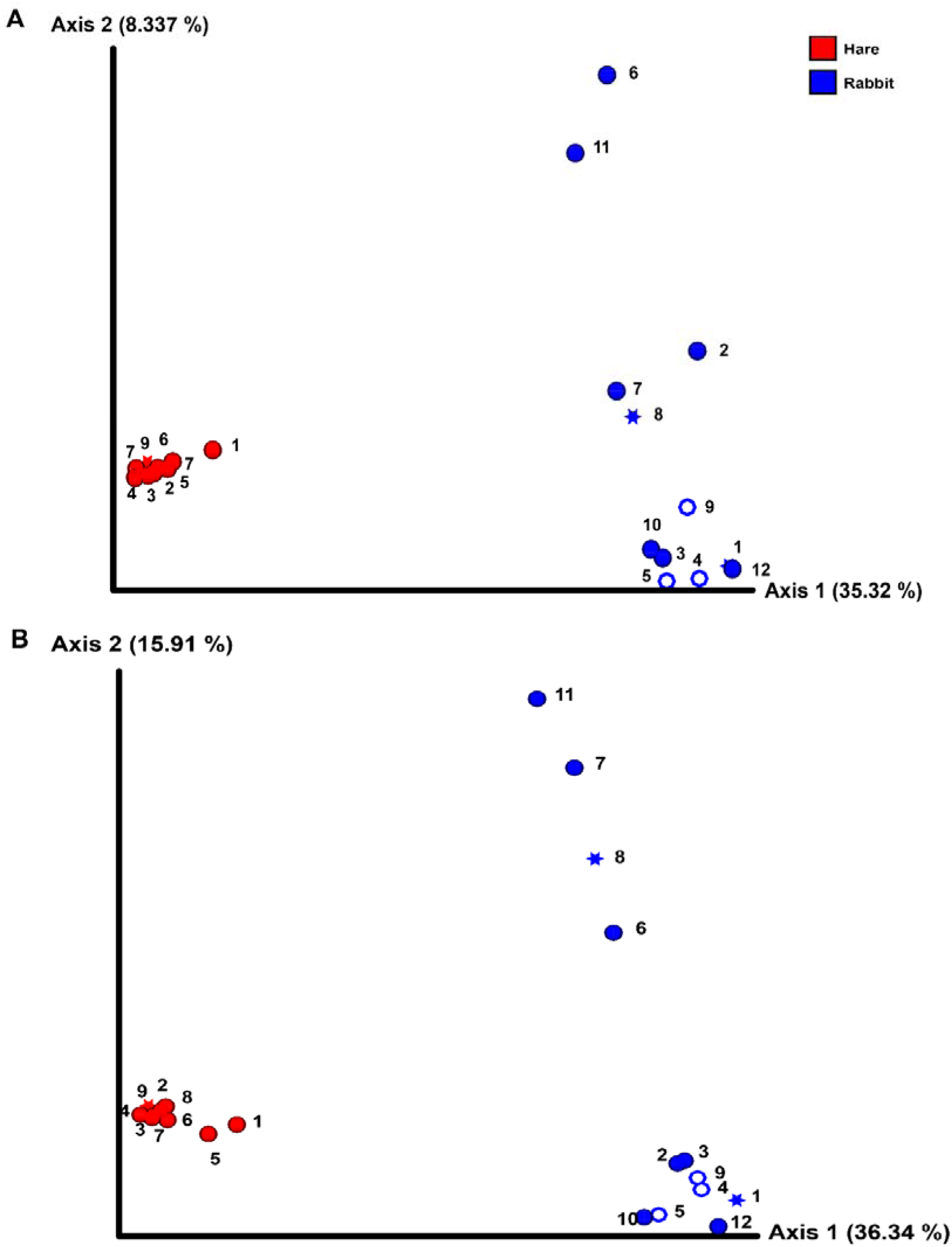
Wild rabbits have greater faecal microbial diversity than sympatric wild hares. 16S rRNA profiling was conducted on wild hare and rabbit faecal samples using either (A) Illumina, or (B) Nanopore sequencing platforms. Microbial species richness and abundance (beta diversity) was estimated using a Bray-Curtis dissimilarity matrix. Significance was assessed using PERMANOVA. Numbers refer to individual animal identifiers as described in Table S1. Stars symbolises pregnant and lactating females, open circles symbolise lactating females, and closed circles symbolise non-pregnant and non-lactating animals.

Taxonomic classification of both the Illumina and Nanopore sequencing data sets identified *Firmicutes* and, to a lesser extent *Bacteroidetes*, to be the two dominant phyla in both hare and rabbit faecal samples (Fig. 2). On average, *Firmicutes* and *Bacteroidetes* together comprised more than 85% of the faecal microbiome in hares and rabbits and across both platforms. Hare faecal samples contained a significantly higher ratio of *Firmicutes* to *Bacteroidetes* compared to rabbit faecal samples (p = 0.015 as measured by Student’s t-test). In both hare and rabbit faecal samples, *Firmicutes* were predominantly represented by the families *Ruminococcaceae*, *Christensenellaceae*, *Lachnospiraceae* (genus *Roseburia*), *Eubacteriaceae*, and *Erysipelotrichaceaea*, while *Bacteroidetes* were predominantly represented by the families *Rikenellaceae* and *Barnesiellaceae* (Fig. 3, Fig. S1). Within the *Firmicutes* and *Bacteroidetes*, the families *Marinifilaceae* (genera *Odoribacter* and *Butryicimonas*), *Tannerellaceae* (genus *Parabacteroides*), and *Veillonellaceae* were more abundant in hares compared to rabbits, while the families *Clostridiaceae 1*, *Enterococcaceae*, *Planococcaceae*, *Lactobacillaceae*, *Peptostreptococcaceae*, and *Bacillaceae* were more abundant in rabbits, although not all families were present in all rabbits (Fig. 3, Fig. S1). Phylum *Tenericutes* (genus *Anaeroplasmataceae*) was present at a significantly higher relative frequency in rabbit faecal samples (p < 0.01 as measured by Student’s t-test). Phylum *Verrucomicrobia* (genus *Akkermansia*) was identified only in rabbit samples and not in hare samples, while phyla *Lentisphaerae* (genus *Victivallis*) and *Synergistetes* (genera *Pyramidobacter*) were identified only in hare samples. Other phyla such as *Proteobacteria* (genus *Sutterella* in both hares and rabbits, family *Desulfovibrionaceae* in hares only, and genus *Enterobacter* in rabbits only), *Actinobacteria* (genus *Eggerthellaceae*), and *Patescibacteria* (family *Saccharimonadaceae*) were present at low abundance (0.5-3%) in both rabbit and hare samples (Fig. 2, Fig. 3, Fig. S1).

**Figure 2:**
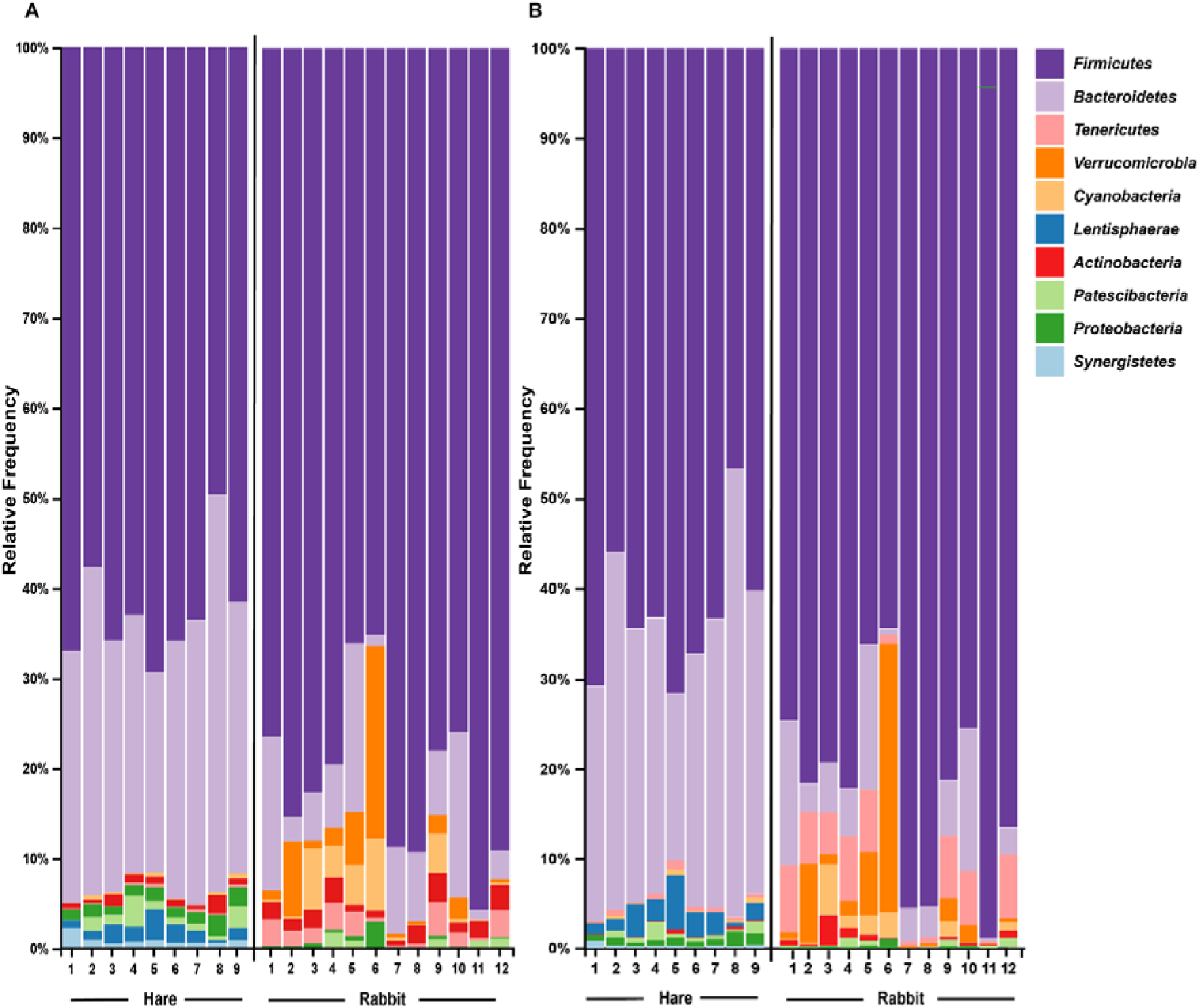
The faecal microbiomes of wild rabbits and hares were distinct at phylum level. Taxonomic classification of 16S rRNA sequences from (A) Illumina, and (B) Nanopore sequencing platforms was performed using BLASTn against the SILVA_132 reference database. The relative frequency of reads assigned to bacterial phyla present at an abundance greater than 0.5% were plotted for each sample.

**Figure 3:**
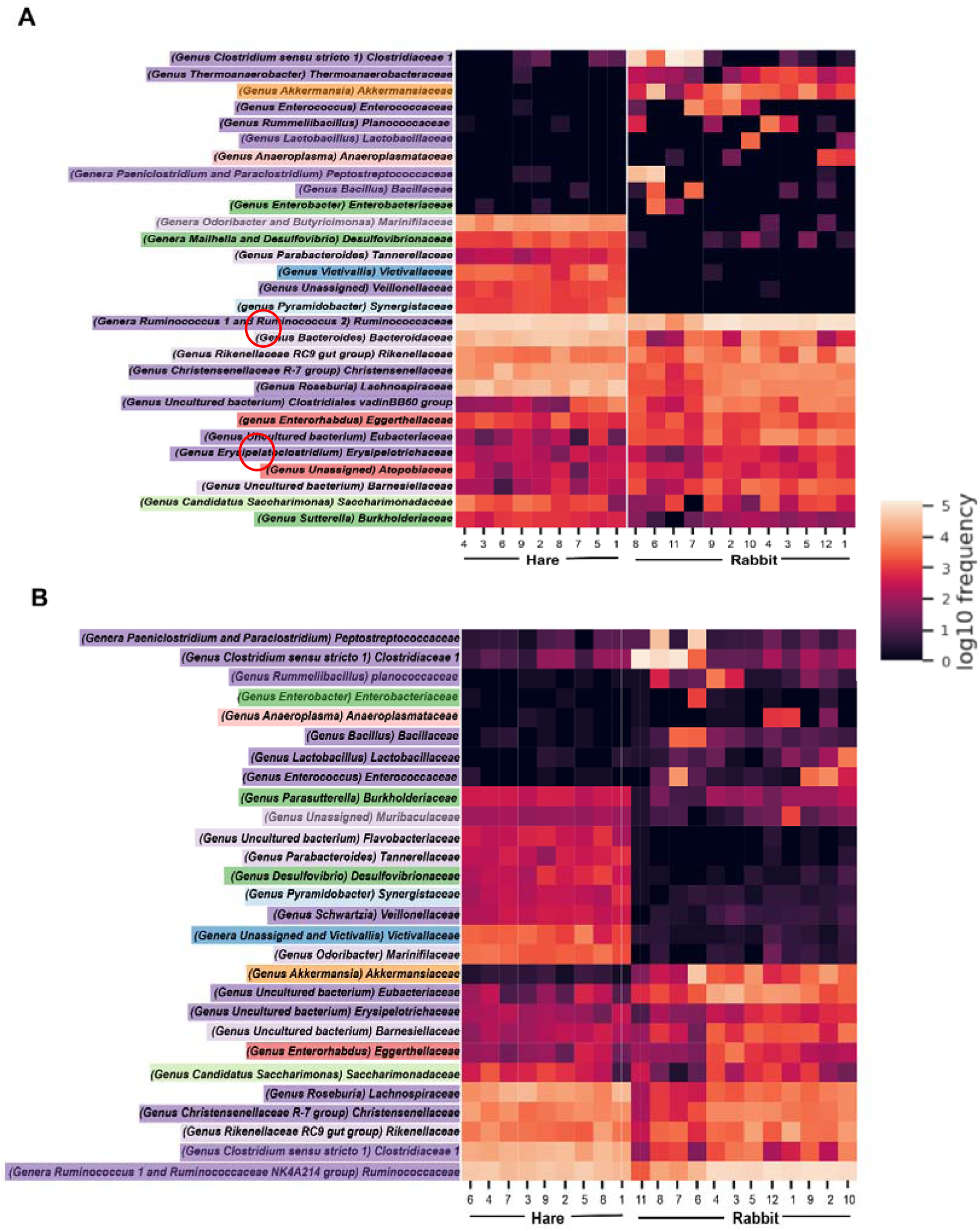
Faecal microbial diversity of Australian wild hares and rabbits at the family level identified unique profiles for wild hares and rabbits. Taxonomic classification of 16S rRNA sequences from (A) Illumina, and (B) Nanopore platforms was performed using BLASTn against the SILVA_132 reference database. The frequency of reads mapping to bacterial families for individual samples are shown in a heatmap on a log10 scale. Family and genera names are highlighted according to phylum. Rows are clustered based on features identified across all samples. Bacterial families present at a frequency less than 0.5% are not included.

The NTC microbiome comprised the phyla *Proteobacteria* and *Bacteroidetes* at relative frequencies of 99% and 1%, respectively (Fig. S2). When analysing the ROCs, there was obvious genomic DNA contamination of both ROCs, with a number of phyla shared between faecal samples and ROCs (Fig. S2). Additional bacterial phyla were also detected only in ROCs, likely comprising the “reagent microbiome”, including *Plantomycetes*, *Dependentiae*, *Chloroflexi*, and *Chlamydiae*.

### Geography alters the faecal microbiome of wild hares in native and introduced region

We then compared the faecal microbiome of Australian wild hares to that of wild hares in their native range of Europe [24] to investigate the influence of geography on the gastrointestinal microbiome. Strikingly, phylum *Spirochaetes* was the third most abundant phyla in faecal samples of European origin, yet it was completely absent from Australian hares and rabbits (Fig. 4). Furthermore, phylum *Patescibacteria* was below our limit of detection (<0.5% relative abundance) in European hare samples, while in Australian hare samples it was present at a higher relative abundance (1.3%). Other phyla were detected in both European and Australian hare samples at similar relative abundances (Fig. 4).

**Figure 4:**
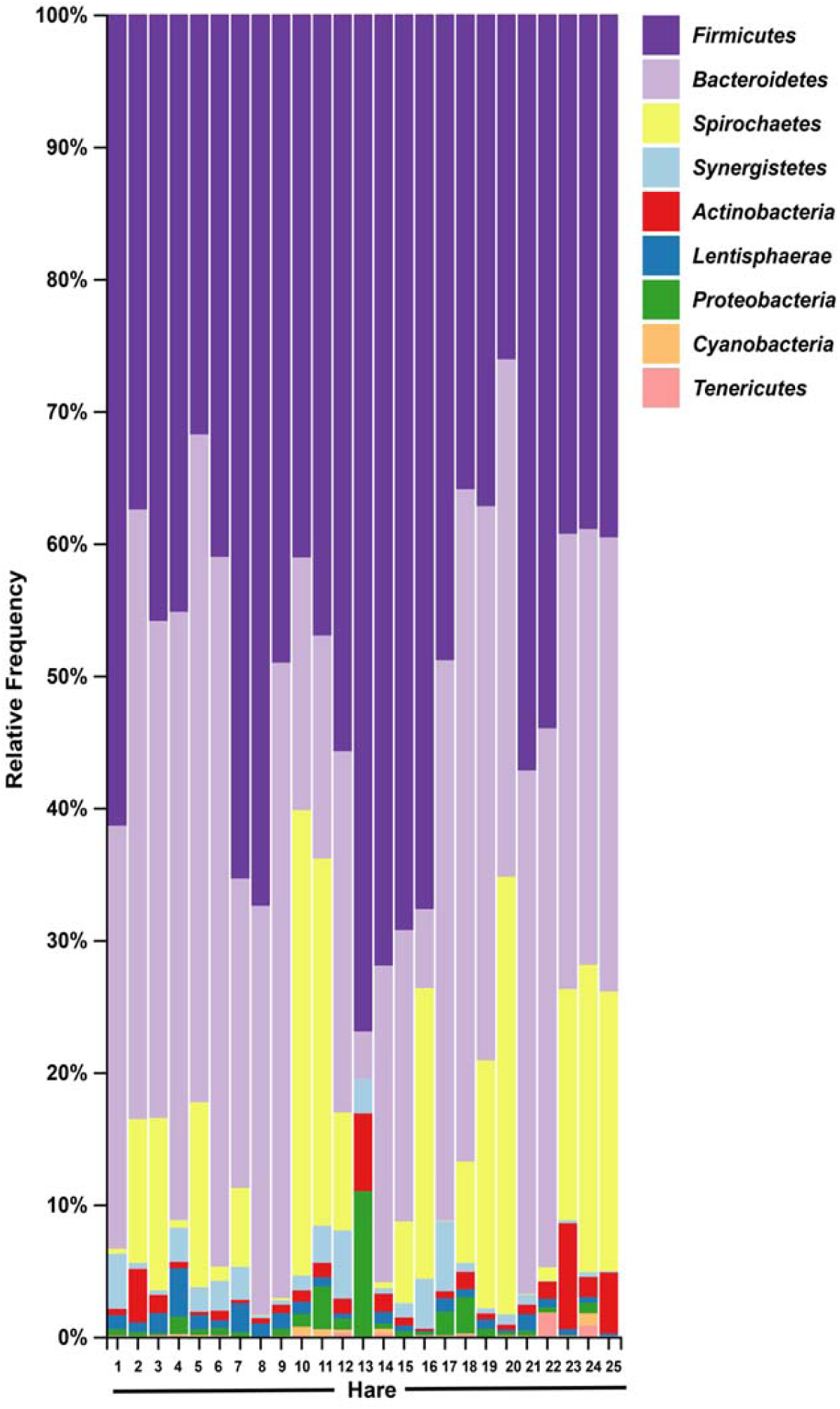
Faecal microbial diversity of wild hares in their native range of Europe shows *Spirochetes* to be the third most abundant bacterial phyla. Raw Illumina sequencing data were obtained from a previous study examining the 16S rRNA diversity in faecal pellets of European brown hares in Europe (24). Reads were processed and classified in parallel with sequencing data from Australia hare faecal samples, using BLASTn against the SILVA_132 reference database. The relative frequency of reads assigned to bacterial phyla present at an abundance greater than 0.5% were plotted for each sample.

### Nanopore and Illumina sequencing platforms reveal a similar faecal microbiome independent of platform

We observed similar faecal bacterial diversity at the phylum, family, and genus levels in both the Illumina and Nanopore datasets, although the relative abundances varied slightly (Fig. 2, Fig. 3, Fig. S1). Most notably, phyla *Actinobacteria* and *Synergistetes* were present at a higher relative abundance in the Illumina dataset, while phyla *Tenericutes*, *Lentisphaerae*, and to a lesser extent *Cyanobacteria*, were present at higher abundance in the Nanopore dataset (Fig. 2). At the family level, *Thermoanaerobacteraceae*, *Bacteroidaceae*, *Clostridiales vadinBB60* group, and *Atopobiaceae* were detected in the Illumina but not the Nanopore data, while *Muribaculaceae* and *Flavobacteriaceae* were identified only in the Nanopore data (Fig. 3, Fig. S1). Even at the genus level there was generally good agreement between datasets (Fig. 3). However, genera *Butyricimonas*, *Mailhella*, *Ruminococcus 2*, *Erysipelatoclostridium*, and *Sutterella* were only detected in Illumina data and *Parasutterella*, *Schwartzia*, an unassigned *Victivallaceae*, and the *Ruminococcaceae NK4A214* group were only identified in the Nanopore data.

## Discussion

To date, studies investigating the gastrointestinal microbial diversity of lagomorphs have been limited to domestic production rabbits in either Europe or China (14-17, 19, 23). Currently only one study has focussed on wild lagomorphs, namely European brown hares in their native European home-range (24). We were interested in understanding why rabbits, an invasive pest species in Australia, were able to rapidly colonise over two thirds of the continent, an area 25 times the size of Britain, within 50 years. In contrast, populations of European brown hares, also a non-native species, remained relatively stable, despite both species occupying a similar ecological niche. Differences in colonising potential of these two species are likely multifactorial, for example, involving differences in behaviour, physiology, and dietary preferences. Comparative analysis of the gastrointestinal microbiome of these two species is one method that can be used to investigate host-specific dietary preferences, which may in turn reveal clues to the differences in environmental impacts that each species has. A secondary aim was to use this study as an opportunity to compare short-read (Illumina) and long-read (Nanopore) sequencing technologies for microbial diversity investigations.

We estimated the gastrointestinal microbiome from faecal samples collected from nine wild European brown hares and twelve European rabbits living sympatrically in their non-native range in Australia. Rabbits had a significantly different faecal microbiome compared to hares and showed greater species richness in their faecal microbiomes and more variation between individuals. Species richness is crucial for imparting resilience to microbial communities, facilitating rapid and effective adaptation to new environmental conditions (47). Species-rich communities can better resist invading pathogens and decreased microbial diversity in humans has been linked to a range of pathologies (47). Hares are known to be more selective foragers than rabbits, which are considered more generalist (7). The observed diversity in the faecal microbiome of wild rabbits may be a consequence of these indiscriminate dietary preferences, or alternatively, a more diverse faecal microbiome may indeed permit rabbits to consume feeds that hares cannot digest. It is interesting to speculate that the high faecal diversity observed in rabbits may allow them to rapidly adapt to new environments, contributing to this species being such a successful invader.

The faecal microbiomes of both host species were dominated by the phyla *Firmicutes* and *Bacteroidetes*, as is typical for other vertebrate species (48). However, the ratio of *Firmicutes* to *Bacteroidetes* varied, being markedly higher in rabbits compared to hares. In humans, an increased *Firmicutes* to *Bacteroidetes* ratio has notoriously been associated with age, diet, and obesity (49). A higher abundance of *Bacteroidetes* (genus *Bacteroides*) has been linked to consumption of diets rich in protein and fat in humans (50), and indeed, hares are known to prefer diets rich in crude fat and protein (51). Despite having a higher daily digestible nitrogen intake, hares tend to have less efficient protein digestion compared to rabbits, potentially due to the absence of key microbes in their gastrointestinal tract (52). Another apparent difference in the faecal microbiomes of these species was the presence of phylum *Verrumicrobia* in rabbits. This phylum is dominated by bacteria from the genus *Akkermansia,* which are noted to be positively influenced by dietary polyphenols (53, 54). Polyphenols are metabolised by intestinal bacteria to generate short-chain fatty acids (SCFAs), and previous studies have demonstrated that rabbits produce a higher concentration of SCFAs than hares, with a higher ratio of butyrate to propionate (55–57). Again, it is unknown whether the *Akkermansia* detected in rabbits in this study permit the digestion of these polyphenols, influencing the dietary preference of rabbits, or whether the intake of polyphenols supports a detectable *Akkermansia* population in rabbits.

Phyla *Lentisphaerae* and *Synergistetes* were observed only in hare populations and were dominated by the genera *Victivallis* and *Pyramidobacter*, respectively. Age and diet appear to be important factors in regulating the abundance of these phyla (58, 59). A previous study in pigs reported a decrease in relative abundance of phyla *Lentisphaerae* and *Synergistetes* with aging (58). Furthermore, lower relative abundances of phyla *Proteobacteria*, *Lentisphaerae*, and *Tenericutes* in vervets and humans have been associated with “Western diets” rich in carbohydrates (59). The observed differences in faecal microbiota could also be related to other known differences in digestive physiology between rabbits and hares. For example, hares have a higher gastrointestinal passage rate compared to rabbits, while rabbits retain digesta longer in order to maximise the efficiency of nutrient extraction (52). Rabbits also have a greater ability to digest hemicelluloses and have a higher rate of methanogenesis compared to hares (55–57, 60).

We also analysed raw data previously sequenced from faecal pellets of wild hares in their native range of Europe to estimate the influence of geography on the gut microbiome (24). We observed the phylum *Spirochaetes* to be the third dominant phyla in the European dataset, in accordance with the original analysis. However, in our Australian dataset only very few reads (less than four) were associated with this phylum. Although bacteria within this phylum can be pathogenic, the *Spirochetes* identified in European hares were associated with a non-pathogenic genus. The absence of this phylum of bacteria in Australian hares may reflect geographical differences in diet between the populations studied, loss of this bacterial species after introduction into Australia, or absence in the original founding hare population. Despite these spirochaetes likely being non-pathogenic, it is worth nothing that the absence of pathogens of invasive species in their non-native range may contribute to spread and persistence of these alien species in new environments. Since faecal samples in this study were collected from healthy hares and rabbits, no candidate pathogens were investigated here.

An additional aim of this study was to compare the bacterial diversity between two emerging sequencing platforms. The Illumina platform is a popular approach for 16S rRNA sequencing, particularly because of its lower error rate (61). However, the short-read length can make species level identification very challenging, especially between closely related species (61). In contrast, the Oxford Nanopore platform has the ability to sequence very long reads, however, it is prone to a relatively high error rate, again making accurate taxonomic assignment challenging (61). In this study, we observed very similar bacterial diversity at the phylum, family, and genus levels with both sequencing platforms, confirming the suitability of either technology for 16S rRNA studies and further confirming the biological relevance of our findings. The minor observed differences in relative abundances of different phyla are most likely due to PCR-based errors or bias, since different primer sets were used for each sequencing platform, or bias during sequencing (61).

In conclusion, we observed notable differences in the microbiome of hares and rabbits living in sympatry in their non-native range in Australia. Though these species inhabit the same habitat, their behaviour and dietary preferences clearly influence differences in faecal bacterial diversity. The more diverse and variable gastrointestinal microbiota of rabbits compared to hares could be a contributing factor in their ability to spread very successfully and establish in new environments. This study also provides additional evidence that the environmental impacts of rabbits are more severe than that of hares, as demonstrated by their less discriminate dietary preferences. Additionally, the absence of bacteria from the phylum *Spirochaetes* in Australian compared to European wild hares demonstrates considerable geographical differences between populations, although whether these spirochetes are beneficial or detrimental to hares was not determined. Future studies correlating different bacterial species in lagomorph microbiomes with particular plant species may provide further insights into the impacts of wild rabbits and hares in Australia.

## Data availability

Detailed protocols are available at https://www.protocols.io/researchers/somasundhari-shanmuganandam/publications. All scripts used were deposited at https://github.com/SomaAnand/Hare_rabbit_microbiome. Raw Illumina sequence data were deposited in the sequence read archive of NCBI under accession number PRJNA576096 and Nanopore data were deposited under accession number PRJNA576096.

---

**Table S1:** Metadata for hare and rabbit samples used in this study.

| **Sample** | **Sample name** | **Sex\*** | **Weight (g)^** | **Collection month** | **Lactation status** |
| --- | --- | --- | --- | --- | --- |
| Hare 1 | MF-136 | NA | NA | January | No |
| Hare 2 | MF-148 | F | 3100 | May | No |
| Hare 3 | MF-149 | M | 3700 | May | No |
| Hare 4 | MF-150 | M | 3040 | June | No |
| Hare 5 | MF-151 | M | 2920 | June | No |
| Hare 6 | MF-152 | F | 3460 | June | No |
| Hare 7 | MF-155 | F | 3250 | July | No |
| Hare 8 | MF-156 | F | 3350 | July | No |
| Hare 9 | MF-157 | F | 4900 | September | Pregnant, lactating |
| Rabbit 1 | MF-138 | F | NA | February | Pregnant, lactating |
| Rabbit 2 | MF-139 | M | 1080 | March | No |
| Rabbit 3 | MF-140 | F | 960 | March | No |
| Rabbit 4 | MF-141 | F | 1380 | March | Yes |
| Rabbit 5 | MF-142 | F | 1440 | March | Yes |
| Rabbit 6 | MF-143 | NA | 400^ | March | No |
| Rabbit 7 | MF-144 | M | 1400 | March | No |
| Rabbit 8 | MF-145 | F | 1500 | March | Pregnant, lactating |
| Rabbit 9 | MF-146 | F | NA | April | Pregnant, lactating |
| Rabbit 10 | MF-147 | F | NA | April | No |
| Rabbit 11 | MF-153 | M | 1500 | June | No |
| Rabbit 12 | MF-154 | M | 1400 | June | No |

\* F female; M male; NA not recorded
^ young rabbit (<12 weeks old)

**Supplementary Figure 1:**
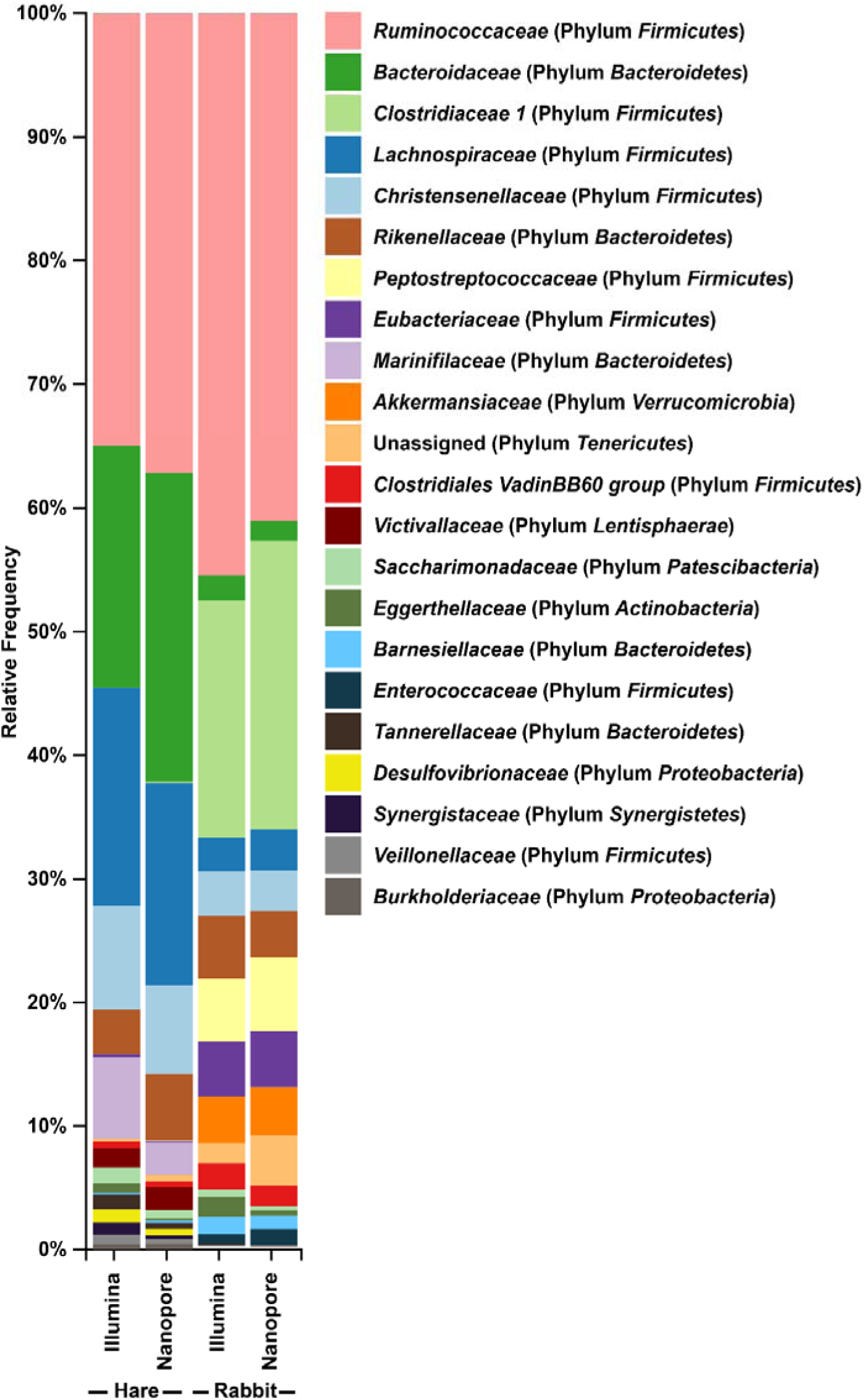
Illumina and Nanopore sequencing platforms produced very similar distributions of bacterial families in wild hare and rabbit faecal samples. Taxonomic classification of 16S rRNA sequences from (A) Illumina, and (B) Nanopore sequencing platforms was performed using BLASTn against the SILVA_132 reference database. Bacterial families present at a relative frequency less than 0.5% are not included.

**Supplementary Figure 2:**
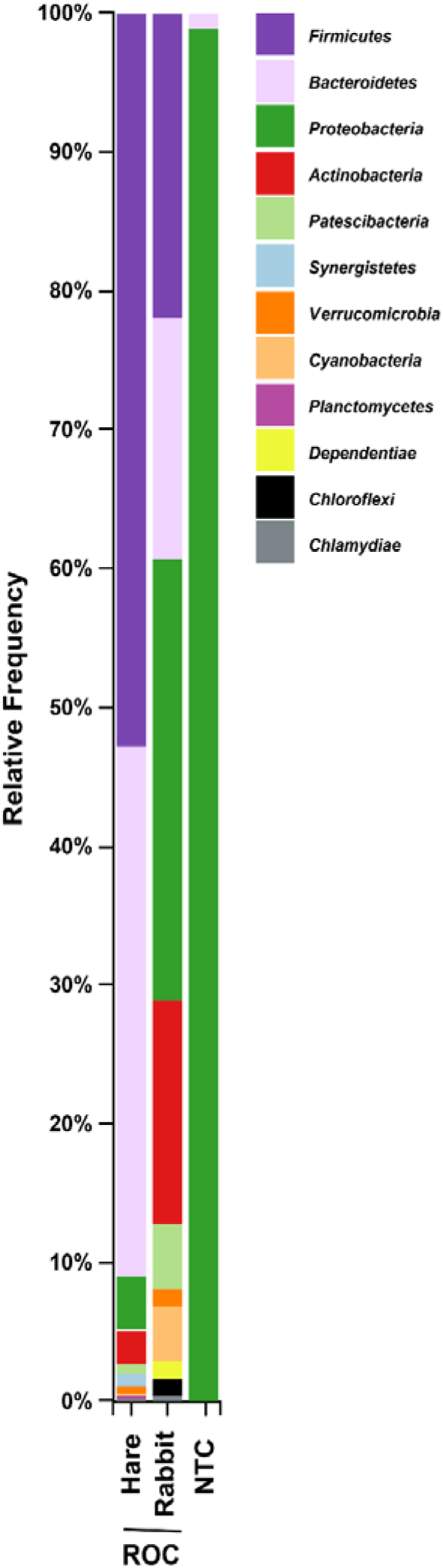
Taxonomic bar plot at phyla rank showing bacterial composition across ROC and NTC samples. Reagent-only controls (ROCs) were extracted and processed in parallel with genomic DNA from faecal pellets. The no template control (NTC) was included in 16S rRNA PCRs and subsequently processed in parallel with samples. Samples were sequenced using the Illumina platform. Taxonomic classification of reads was performed using BLASTn against the SILVA_132 reference database. Bacterial phyla present at a relative frequency less than 0.5% are not included.

